# Genetic Portrait of North-West Indian Population based on X Chromosome *Alu* Insertion Markers

**DOI:** 10.1101/761692

**Authors:** Gagandeep Singh, Yellapu Srinivas

**Affiliations:** Panjab University, Chandigarh; Wildlife Institute of India

**Keywords:** X chromosome, *Alu* insertions, population genetics, polymorphism, human forensics

## Abstract

*Alu* insertion elements represent the largest family of Short Interspersed Nuclear Elements (SINEs) in the human genome. Polymorphic *Alu* elements are stable and conservative markers that can potentially be applied in studying human origin and relationships as they are identical by descent and known for lack of insertion in ancestral state. In this study, 10 *Alu* insertions of X chromosome were utilized to tabulate allele frequency distributions and compute parameters of forensic relevance in the 379 unrelated healthy individuals belonging to four different ethnic groups (Brahmin, Khatri, Jat Sikh and Scheduled Caste) of North-West India. Furthermore, the *D*_A_ and *F*_ST_ values of pairwise interpopulation differentiations, multidimensional scaling and Bayesian structure clustering analysis were also computed to probe the genetic relationships between present studied populations and with other 21 reference populations. Six X-*Alu* insertions were observed to be polymorphic in all the populations, whereas the others appeared as monomorphic in at least one studied population. The insertion allele frequencies were in the range of 0.15 at Ya5DP3 to 0.9888 at Ya5DP77. Most polymorphic *Alu* elements showed moderate to low genetic diversity. The maximum value of power of exclusion (PE) was 0.1645 at Ya5NBC37 marker, whereas the minimum was 0.0001 at Ya5DP4 locus, implying the significance of X chromosome *Alu* elements in forensic genetic investigations. Genetic relationships agree with a geographical pattern of differentiation among populations. The results of present study establish that X chromosome *Alu* elements comprise a reliable set of genetic markers useful to describe human population relationships and structure.

## Introduction

The X chromosome provides a valuable source of information for population genetics and serves as a unique tool in forensic studies as it integrates the desirable features of other commonly used genetic markers. It recombines and has a deep history, similar to the autosomes; and it has accessible haplotypes along with sex-biased mode of inheritance, similar to mtDNA and the Y chromosome (Hamosh et al. 2000; Schaffner et al. 2004). The X chromosome offers many unique features concerning its transmission pattern. Males inherit a single maternal X chromosome, whereas females inherit an X chromosome from each parent. Compared to autosomes, the X chromosome has a lower mutation rate, lower recombination rate and smaller effective population size (Ne), resulting in quicker genetic drift. Such facts contribute to the lower genetic diversity (estimated to be about half of that on the autosomes), and a stronger linkage disequilibrium (LD) and population structure in comparison to autosomes. On the other hand, since X chromosome spends two-thirds of its lifetime in female subjects, its polymorphisms mainly reflect the history of females (Schaffner et al. 2004; Athanasiadis et al. 2007; Ferragut et al. 2017).

The 150 million base pair (Mb) size of the human X chromosome is consistently ∼5% of the genome among mammals (Schaffner et al. 2004). It contains multiple types of non-coding markers distributed along its sequence, including *Alu* insertions, insertion-deletions (INDELs), short tandem repeats (STRs) and single nucleotide polymorphisms (SNPs). *Alu* insertion elements represent the largest family of Short Interspersed Nuclear Elements (SINEs) in humans and are classified into 12 subfamilies that appeared at different times during primate evolution (Schmid and Deininger 1975; Kapitonov and Jurkal 1996). *Alu* elements are dimorphic for the presence or absence of the insertion, have a known ancestral state (lack of insertion), free from homoplasy (identical by descent) and are selectively neutral DNA markers (Batzer et al. 1994; Carroll et al. 2001; Stewart et al. 2011). All of these distinctive attributes make the human *Alu* insertion elements a good tool for studying the genetic variation and the evolutionary relationships of human populations (Stoneking et al. 1997).

In large-scale geographical studies, X chromosome variation provides insight into modern human history indicating that Non-African populations were unlikely to be derived from a very small number of African lineages (Yu et al. 2002). However, in fine-scale geographical studies, the use of X chromosome variation is scarce, when compared with the large number of human population studies focusing on Y chromosome and mtDNA markers. So far, there is not even a single study available in the literature to assess the variability of *Alu* polymorphisms of the X chromosome in the Indo-European speaking population groups from Punjab, North-West India. Due to its geographical location, the present day North-West Indian population may offer unique insights into historic migration events. The exposure of this region’s population with the different ethnic background people from the pan-world might have created a mosaic-like pattern in their genomic ancestry (Ahloowalia 2009). Therefore, for the first time, in this work, a set of 10 X chromosome *Alu* insertions in four Indo-European speaking ethnic groups (Brahmin, Jat Sikh, Khatri and Scheduled Caste) of Punjab, North-West India were analyzed. The objectives were to (1) to explore the extent of genetic variations of polymorphic X chromosome *Alu* insertions in ethnic groups of North-West India, and (2) to apply this variation for studying population relationships among different ethnic groups in a broader geographic context and to evaluate their usefulness in a forensic frame of reference.

## Materials and methods

### DNA samples

Blood samples were collected from 379 unrelated healthy individuals (175 males and 204 females) belonging to four different ethnic groups (Brahmin, Khatri, Jat Sikh and Scheduled Caste) of Punjab, North-West India. Genomic DNA was isolated from blood samples using standard inorganic method with few laboratory modifications (Miller et al. 1988). All samples were collected after obtaining the informed consent of participants and were analyzed anonymously. The work was approved by the ethical committee of the Panjab University, Chandigarh as well as that of Guru Nanak Dev University, Amritsar, Punjab, India. Complete information regarding studied ethnic groups including their geographic origin, ethnicity, pedigree, inclusion and exclusion criteria were given in our earlier study (Singh et al. 2016). All above methods were performed in accordance with the relevant guidelines and regulations.

### Genetic markers and genotyping

All PCR amplifications were carried out in 10 μl reactions for a set of 10 human-specific X chromosome *Alu* insertions (Ya5DP4, Ya5DP3, Ya5DP77, Yb8DP49, Ya5491, Ya5NBC37, Yb8NBC102, Yb8NBC634, Ya5DP62 and Ya5DP13). Primer sequences and amplification conditions were obtained from previous studies (Callinan et al. 2003; Di Santo et al. 2018). The efficiency and reliability of the PCR reactions were monitored using control reactions. Genotypes were identified by 2-3% agarose electrophoresis of PCR products and visualized under UV light in the presence of ethidium bromide dye.

### Statistical analyses

Allele frequencies were calculated from the genotype data by direct gene counting method at each locus separately for each ethnic group (Li 1976). Heterozygosity and exact test of Hardy- Weinberg equilibrium (HWE) for female samples, pairwise exact test of linkage disequilibrium (LD) for male samples and *F*_ST_ values for both males and females were calculated using Arlequin software v3.5 (Excoffier and Lischer 2010). Statistical parameters of forensic relevance were estimated using online software X-chromosome STR homepage (http://www.chrx-str.org) (Szibor et al. 2006). The Bayesian clustering analysis implemented in the program STRUCTURE v2.3 (Pritchard et al. 2000) was used to detect population structure assuming population admixture model and correlated allele frequencies within the population. All runs were performed using a burn-in period of 10^5^ iterations followed by 500,000 iterations of MCMC repeats. Value K was allowed to vary between 1 and 6 with 10 independent runs for each K value. To choose the optimal value of K, web-based program Structure Harvester was used. Finally, all the graphic displays were obtained using the Pophelper web page (Francis 2017). In order to examine the relationship of the present studied groups with other published data, pairwise Nei genetic distance (*D*_*A*_) and fixation index (*F*_ST_) were calculated from allele frequency data with the help of software POPTREE2 (Takezaki et al. 2009). The heat maps were constructed based on *F*_ST_ and *D*_*A*_ values using *R* statistical software v3.3. For easier visualization of the genetic distances, a multidimensional scaling (MDS) plot of the pairwise *F*_ST_ values was represented using SPSS v16.0.

## Results and discussion

### Genetic diversity and forensic efficiency parameters

Allele frequencies for the 10 *Alu* insertions in four studied populations are depicted in **Fig. 1.** Six X *Alu* insertions (Ya5DP3, Yb8DP49, Ya5NBC37, Yb8NBC102, Yb8NBC634, Ya5DP62) were observed to be polymorphic in all the populations, whereas the others appeared as monomorphic in at least one studied population (Ya5DP4 in Khatri and Scheduled Caste, Ya5DP13 in all the populations for the absence of insertion; and Ya5DP77 in Brahmin and Ya5491 in all the populations for the insertion). The insertion allele frequencies across the studied populations were in the range of 0.15 at Ya5DP3 to 0.9888 at Ya5DP77. Significant deviations from HWE were found in few *Alu* insertion polymorphisms in female samples and on the other hand no significant linkage disequilibrium was present in any pair of the *Alu* markers in male samples. In general, the allele frequencies found range within the general patterns as described in previous studies (Athanasiadis et al. 2007; Ferragut et al. 2017; Di Santo et al. 2018). In the present study, similar to many other studies, values of minor allele frequency (MAF) of some loci were generally low, further indicating that X chromosome *Alu* elements have high potential to detect population structure and to study relationship between populations (Da la and Raska 2014). Most polymorphic *Alu* elements showed moderate to low genetic diversity. **Table 1** shows the gene diversity and some parameters of forensic relevance calculated for each marker and population. Ya5NBC37 displayed the highest heterozygosity (0.3895) and Ya5DP4 the lowest (0.0263). Likewise, the most diverse population seems to be the Khatri (H= 0.1087) followed by the Scheduled Caste (0.0891), Jat Sikh (0.0571) and Brahmin (0.0509). The average gene diversity value of X chromosome *Alu* insertions in the present study was 0.027 similar to that observed in other European and Asian populations (Ferragut et al. 2017; Di Santo et al. 2018). The value of polymorphic information content (PIC) was in the range of 0.0104 to 0.3608 with a mean value of 0.1521. Additionally, the maximum value of power of exclusion (PE) was 0.1645 at Ya5NBC37 marker, whereas the minimum was 0.0001 at Ya5DP4 locus. Overall values obtained for power of discrimination were moderate in both females and males. Therefore, the present study further provides and highlights the importance of X chromosome markers for forensic casework and personal identification.

**Table 1.**
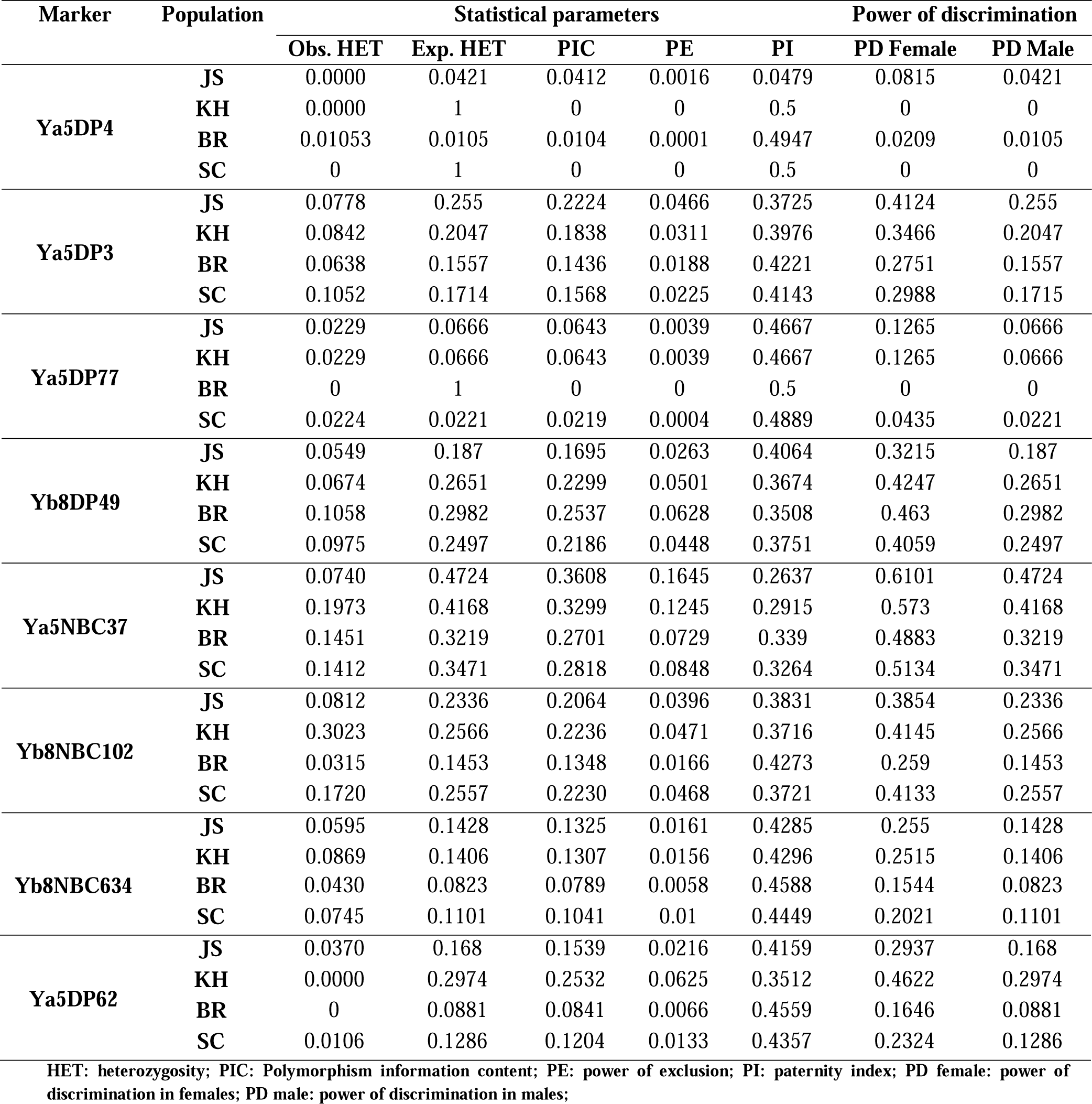
Gene diversity and forensic parameters of X chromosome *Alu* markers (Jat Sikh: 94; Khatri: 93; Brahmin: 80 and Scheduled Caste: 70)

**Figure 1.**
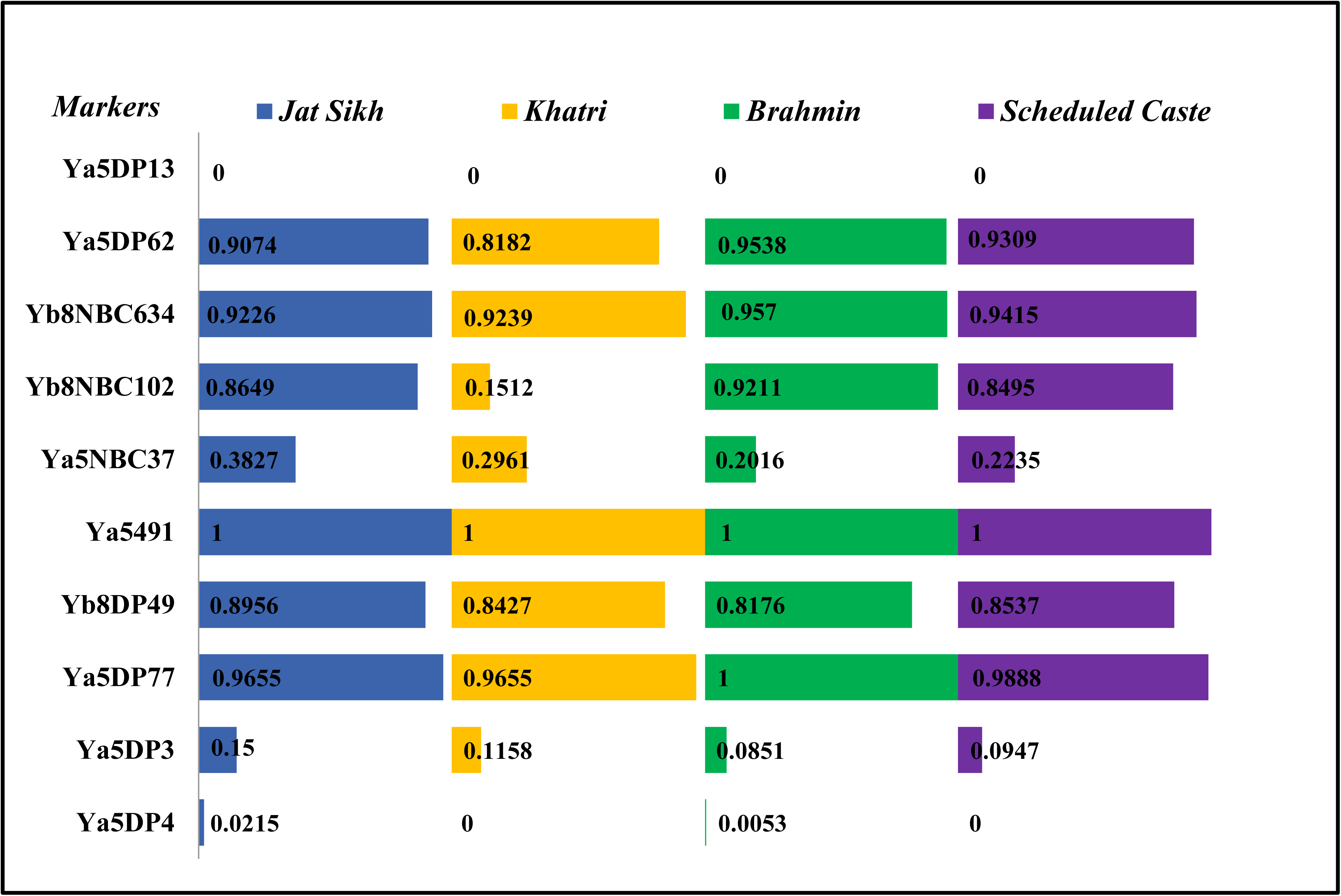
Plots of insertion allele frequencies of 10 X *Alu* markers (Jat Sikh: 94; Khatri: 93; Brahmin: 80 and Scheduled Caste: 70)

The *F*_ST_ values of the *Alu* insertions for the populations analysed in this study were in the range of 0-0.19 with no significant differences from each other, reflecting that they share, at least partially a common ancestry or gene pool (**Supplementary Table 1**). The absence of marked interpopulation differentiation observed in this study is in accordance with the lower mutation rate in *Alu* insertions as compared to the other markers, and therefore, helpful in gaining insights into population structure and relationships between the populations (Athanasiadis et al. 2007; Di Santo et al. 2018; Gayà□vidal et al. 2010).

### Genetic structure analysis

In the present study, Bayesian clustering analysis in STRUCTURE v2.3 (Pritchard et al. 2000) was performed to reflect the memberships of biogeographic ancestry components for the present studied populations. The Bayesian analysis suggested the presence of two genetic clusters (K=2) in the studied populations. As shown in **Fig. 2**, the four populations (Jat Sikh, Brahmin, Khatri and Schedule Caste) are separated by white lines and each single vertical line represents one individual which was partitioned into different coloured segments based on individual estimated membership fractions. At K=2, the Khatri population was separated from remaining three populations with some admixture from other populations. Similarly with an increase in K (K=2- 4), as shown in **Fig. 2**, the studied populations shared mixed membership in different colour components with no sub structuring in the studied groups. The results are in accordance with the previous studies on the ethnic groups of North-West Indian populations using different sets of molecular markers (Singh et al. 2017; Singh et al. 2019). The results further shred evidence and points out that the studied populations from North-West region, India might have reconstituted themselves a number of times in the wake of innumerable foreign invasions and migrations over the centuries (Guha, 1931).

**Figure 2.**
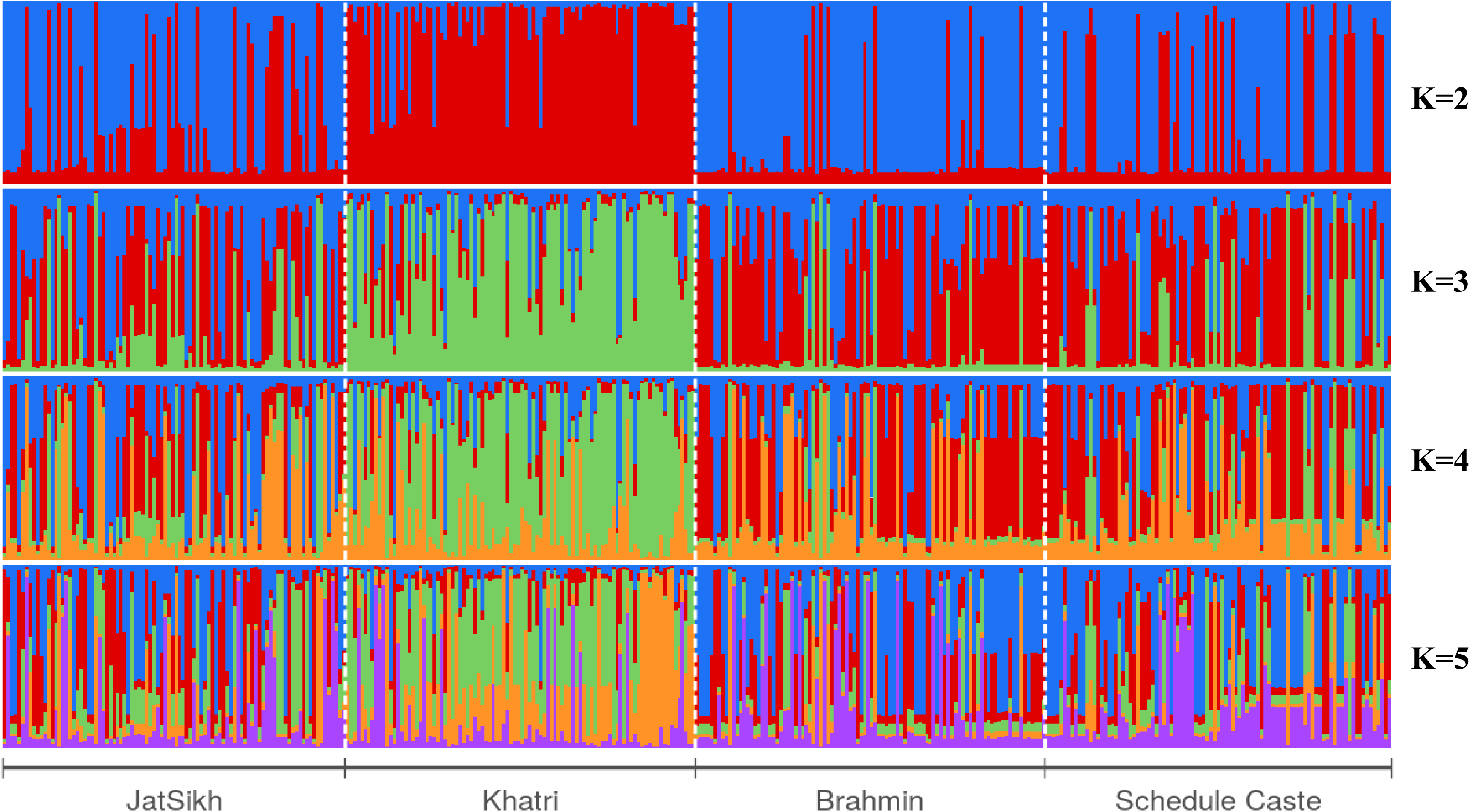
Population Structure analysis of 4 studied populations at K=2 to K= 5

### Comparison with other populations

To explore the genetic differences, the studied populations were compared with previously published data on populations from Asia, America, Europe and Africa (Athanasiadis et al. 2007; Ferragut et al. 2017; Di Santo et al. 2018; Gayà□vidal et al. 2010; Ferragut et al. 2018; Resano et al. 2016) using pairwise *F*_ST_ and Nei’s *D*_A_ genetic distance measures. The *F*_ST_ is generally considered as a measure of population differentiation on account of genetic structure (Jakobsson et al. 2013) and is directly related to the variance in allele frequency among populations. The larger *F*_ST_ value, the higher genetic differentiation between populations is, and vice versa. *F*_ST_ values observed were in the range of 0.09 to 0.832 with maximum variation findings between 0.4 to 0.8. Similarly, North-West Indian populations used in this study have moderate and equal proportions (*F*ST values 0.5-0.7) of genetic differentiation with other populations elsewhere. As shown in heatmap (**Fig. 3a),** the deeper green colour represents the larger *F*_ST_ value, indicating more genetic differentiation; conversely, the deeper yellow colour represents the smaller *F*_ST_ value with less genetic differentiation between geographically closer populations.

**Figure 3.**
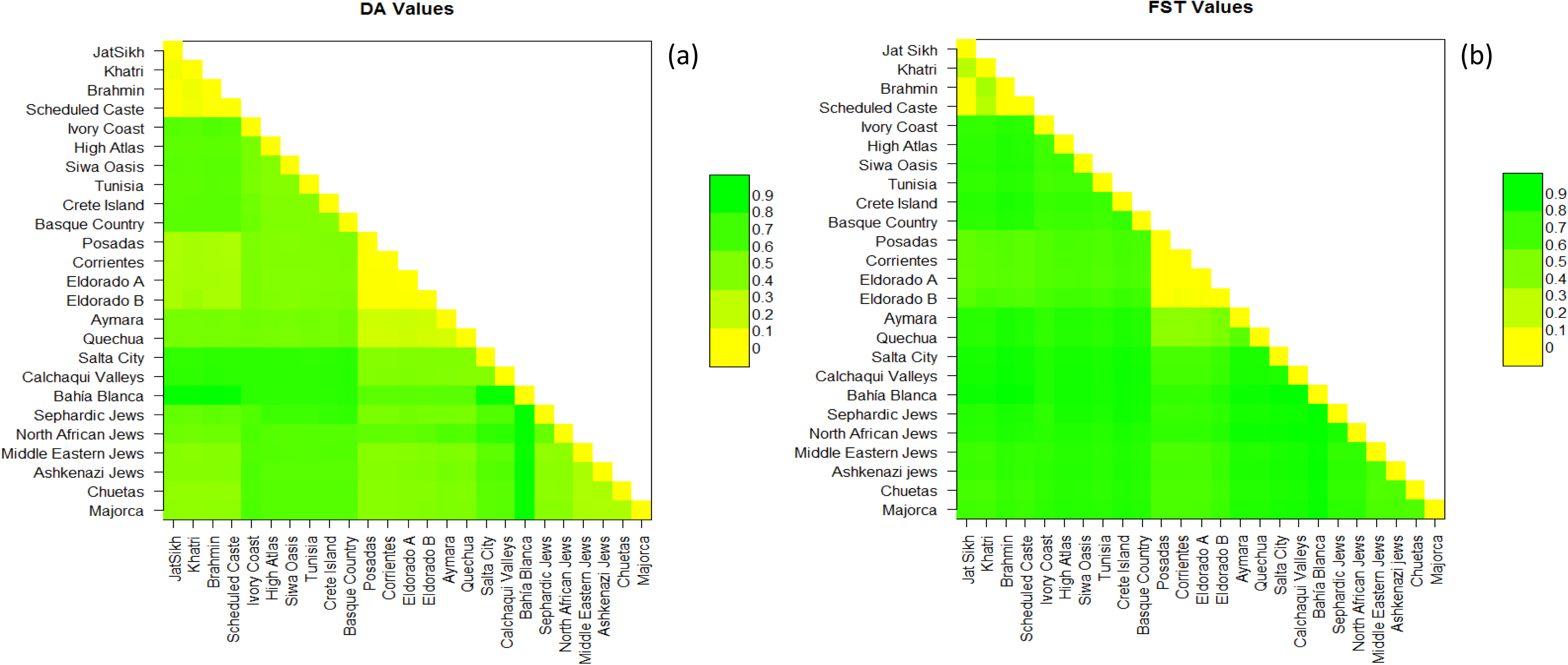
(a) Heat map of pairwise *F*_*ST*_ values of 10 X chromosome *Alu* elements in North- West Indian population and 21 previously studied populations based on *R* software. (b) Heat map of pairwise *D*_*A*_ values of 10 X chromosome *Alu* elements in North-West Indian population and 21 previously studied populations conducted with *R* software.

Nei’s *D*_A_ distance is another most commonly used genetic distance measure to identify the genetic divergence between populations and developed under the assumption that genetic difference originated due to genetic drift and mutation events (Nei and Roychoudhury 1974). Nei’s *D*_A_ distance values were in the range of 0.004 to 0.61, with maximum findings between 0.2 to 0.5. As shown in the heatmap (**Fig. 3b)**, the deeper green colour represents greater *D*_A_ values indicating larger genetic distances, while, the deeper yellow colour represents smaller *D*_A_ values with closeness among populations. Dark coloured blocks were observed between present studied and other different continental populations indicating a greater genetic distance between North- West Indian Populations and reference population used in this study. On the other hand, light coloured blocks were observed between ethnic group of present study, indicating the genetic closeness in comparison to other reference populations indicating genetic closeness to some extent.

Further, to evaluate the genetic relationship between present and other previously studied populations (**Supplementary Table 2**) a multidimensional scaling (MDS) plot was constructed using pairwise *F*_ST_ genetic distance values (**Fig. 4**). All the populations were exhibited with small icons and it was noticeable that the population distribution in the plot was in general concordance with their geographic affiliations. *Alu* insertions of X chromosome clearly discriminate between continents with European, Asian and African populations distantly located on the plot. With the available data, all the Asian populations appeared closer to European and Amerindian populations were placed in between African and European population clusters. Also the four ethnic groups of this study form a closer cluster, suggesting a homogenous genetic entity. These results are in concordance with those obtained in autosomal and uniparental (mtDNA) markers, which showed that those inhabiting in the same geographical terrain show closer genomic affinity irrespective of the social and cultural rankings of the populations (Majumder et al. 1999).

**Figure 4.**
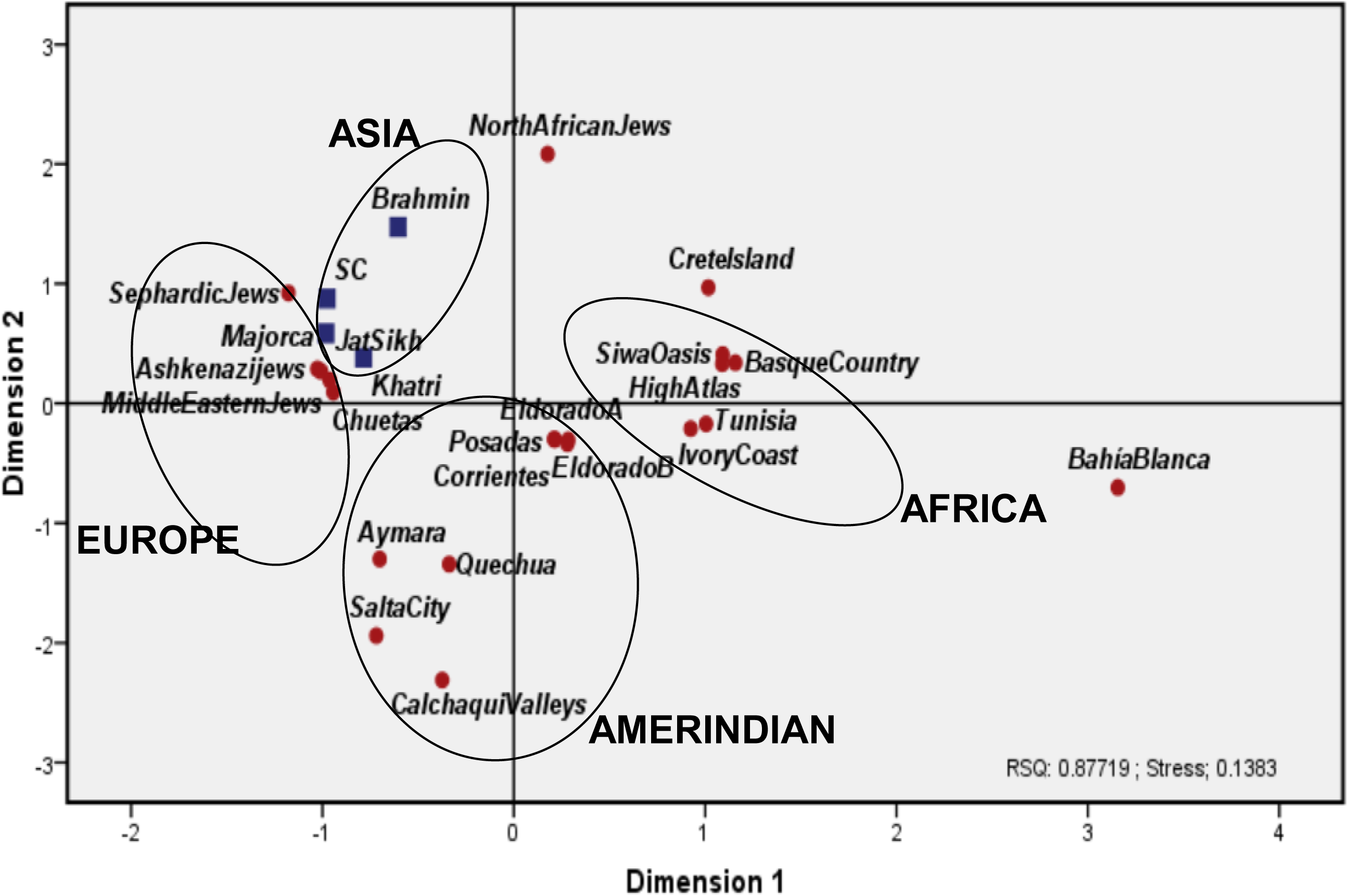
Multidimensional scaling analysis (MDS) based on *F*_*ST*_ genetic distances calculated for 10 X chromosome Alu elements

The genetic analyses of the present study further confirms the importance of using X chromosomal binary markers, such as *Alu* insertions, for differentiating the population of North- West India, where a different level of admixture through migrations could be observed. This study provides a useful database of 10 X chromosome markers for the population of North-West India and highlights the genetic homogeneity among the ethnic groups of Punjab (North-West India). The results obtained from these X chromosome *Alu* elements comprise a reliable set of genetic markers that could assist in forensic genetic investigations. This work also highlights the importance of new studies on additional populations of the Indian subcontinent, as no comparable data exist in literature on X chromosome *Alu* insertions to complete the genetic portrait of the Indian population. Therefore, future investigations focusing on a wider range of X chromosome markers from diverse populations will surely be helpful in some unresolved aspects. We believe the data generated from present study can be meaningful for further enriching the genetic background researches from Indian subcontinent.

## Acknowledgements

We are grateful to all of the donors for providing blood samples and the people who contributed to the collection. Financial support for carrying out this study from DBT through project grant: BT/PR2722/Med/13/121/2001 (DBT, India) to Bhanwer AJS at Guru Nanak Dev University, Amritsar is acknowledged. Additionally, financial support from UGC under Centre with Potential for Excellence in Particular Area (CPEPA) and University with Potential for Excellence (UPE) to AJS Bhanwer at Guru Nanak Dev University, Amritsar is also acknowledged. The authors are specially thankful to Dr. Jim Lemon, for helping with R codes needed for genetic analysis.

