## Supplementary Table 1 for "Genetic Portrait of North-West Indian Population based on X Chromosome *Alu* Insertion Markers"

**Supplementary Table 1 : F_ST_ values for *Alu* markers for all the analyzed populations**

| **Populations** | **F_ST_ values** | **P values** |
| --- | --- | --- |
| Jat Sikh *vs* Khatri | 0.009 | 0.474 |
| Jat Sikh *vs* Brahmin | 0.0199 | 0.058 |
| Jat Sikh *vs* Scheduled Caste | 0.013 | 0.105 |
| Khatri *vs* Brahmin | 0.001 | 0.416 |
| Khatri *vs* Scheduled Caste | 0 | 0.553 |
| Brahmin *vs* Scheduled Caste | 0 | 0.965 |
