## Supplementary Table 2 for "Genetic Portrait of North-West Indian Population based on X Chromosome *Alu* Insertion Markers"

**Supplementary Table 2 :** Samples used in the population comparisons, number of chromosomes (N) studied and respective references

| **Population** | **N** | **Reference** |
| --- | --- | --- |
| Jat Sikh | 94 | Present study |
| Khatri | 93 | Present study |
| Brahmin | 80 | Present study |
| Scheduled Caste | 70 | Present study |
| Sephardic Jews | 103 | Ferragut *et al*., 2017 |
| North African Jews | 77 | Ferragut *et al*., 2017 |
| Middle Eastern Jews | 75 | Ferragut *et al*., 2017 |
| Ashkenazi Jews | 85 | Ferragut *et al*., 2017 |
| Chuetas | 140 | Ferragut *et al*., 2017 |
| Majorca | 89 | Ferragut *et al*., 2017 |
| Ayamara | 152 | Gaya-Vidal *et al*., 2010 |
| Quechua | 147 | Gaya-Vidal *et al*., 2010 |
| High Atlas | 150 | Athanasiadis *et al*., 2007 |
| Siwa Oasis | 143 | Athanasiadis *et al*., 2007 |
| Tunisia | 165 | Athanasiadis *et al*., 2007 |
| Ivory Coast | 73 | Athanasiadis *et al*., 2007 |
| Crete | 119 | Athanasiadis *et al*., 2007 |
| Basque Country | 133 | Athanasiadis *et al*., 2007 |
| Posadas | 52 | Di Santo *et al*., 2018 |
| Corrientes | 92 | Di Santo *et al*., 2018 |
| Eldorado A | 27 | Di Santo *et al*., 2018 |
| Eldorado B | 27 | Di Santo *et al*., 2018 |
| Salta City | 153 | Ferragut *et al*., 2018 |
| Calchaqui Valleys | 121 | Ferragut *et al*., 2018 |
| Bahía Blanca | 62 | Resano *et al*., 2016 |
